# Applying biophysical models to understand the role of white matter in cognitive development

**DOI:** 10.1101/347872

**Authors:** Elizabeth Huber, Rafael Neto Henriques, Julia P. Owen, Ariel Rokem, Jason D. Yeatman

**Author notes:** Correspondence: Elizabeth Huber, Institute for Learning & Brain Sciences, Portage Bay Building, Box 357988, University of Washington Seattle, WA 98195, USA. Declarations of interest: none.

## Abstract

Diffusion MRI (dMRI) holds great promise for illuminating the biological changes that underpin cognitive development. The diffusion of water molecules probes the cellular structure of brain tissue, and biophysical modeling of the diffusion signal can be used to make inferences about specific tissue properties that vary over development or predict cognitive performance. However, applying these models to study development requires that the parameters can be reliably estimated given the constraints of data collection with children. Here we collect repeated scans using a multi-shell diffusion MRI protocol in a group of children (ages 7-12) and use two popular biophysical models to characterize axonal properties. We first assess the scan-rescan reliability of model parameters and show that axon water faction can be reliably estimated from a relatively fast acquisition, without applying spatial smoothing or de-noising. We then investigate developmental changes in the white matter, and individual differences in white matter that correlate with reading skill. Specifically, we test the hypothesis that previously reported correlations between reading skill and diffusion anisotropy in the corpus callosum reflect increased axon density in poor readers. Both models support this interpretation, highlighting the utility of biophysical models for testing specific hypotheses about cognitive development.

## 1. Introduction

White matter biology has been studied extensively using invasive techniques in nonhuman animals (reviewed in (Walhovd, Johansen-Berg, & Karadottir, 2014)). A promising approach for investigating the maturational trajectory of human white matter, and the neurobiological underpinnings of complex cognitive skills like reading, is to leverage models that link biology to non-invasive MRI measurements. Biophysical models assign biological interpretations to metrics derived empirically from diffusion MRI (dMRI) techniques (Nilsson, van Westen, Stahlberg, Sundgren, & Latt, 2013), allowing inferences about the specific white matter properties that underlie individual differences in perception, cognition, and learning. For these biophysical models to have widespread application in developmental cognitive neuroscience, it is important to ascertain whether the parameters derived from these models reliably index individual differences in the white matter of developing children.

Diffusion tensor imaging (DTI) is the most widely used dMRI technique to studying white matter development, owing partially to the fact that the model is easy to fit and highly reliable. DTI has revealed protracted development of the human white matter with development, continuing throughout adolescence and into adulthood (Lebel & Beaulieu, 2011; Mukherjee et al., 2001). However, DTI metrics lack biological specificity. For example, developmental variation in fractional anisotropy (FA) can reflect differences in axonal packing density, caliber, myelination, or spatial coherence, as well as changes in the number, size, and branching of glial cells (Alexander, Lee, Lazar, & Field; Basser & Pierpaoli, 1996; De Santis, Drakesmith, Bells, Assaf, & Jones, 2014; Jeurissen, Leemans, Tournier, Jones, & Sijbers, 2013; Jones, Knosche, & Turner, 2013; Walhovd et al., 2014).

Two popular modeling approaches for estimating biologically specific properties of the white matter from dMRI data are the “White Matter Tract Integrity” model (WMTI (Fieremans, Jensen, & Helpern, 2011; Fieremans, Novikov, Jensen, & Helpern, 2010)) and the “Neurite Orientation Dispersion and Density Imaging” model (NODDI (Zhang, Schneider, Wheeler-Kingshott, & Alexander, 2012)). Although these models makes certain simplifying assumptions (reviewed in (Novikov, Kiselev, & Jespersen, 2018)), they have been successfully applied to the study of development (Jelescu et al., 2015), aging (Benitez et al., 2014 2015), and white matter pathology (Benitez et al., 2014; Falangola et al., 2014; Fieremans et al., 2013; Guglielmetti et al., 2016; Jelescu et al., 2016; Kelm et al., 2016), and both provide estimates of tissue properties that are consistent with histological measurements (reviewed in (Jelescu & Budde, 2017)). Particularly, NODDI has been used to disentangle developmental changes related to dispersion versus density of neurites (axonal and/or dendritic processes) over development (Chang et al., 2015; Genc, Malpas, Holland, Beare, & Silk, 2017; Kodiweera, Alexander, Harezlak, McAllister, & Wu, 2016; Mah, Geeraert, & Lebel, 2017). With the successful application of these models in the context of development, they are now beginning to be applied to the study of individual differences in cognition (Chung et al., 2018). To date, these models have not been applied to the study of reading development.

The relationship between individual differences in reading and white matter diffusion properties has been studied extensively using DTI (Wandell & Yeatman, 2013). While this approach has identified several anatomical correlates of skilled reading, it has often produced conflicting and counterintuitive results. For example, correlations between FA and reading skill are consistently reported in a number of anatomical tracts (Ben-Shachar, Dougherty, & Wandell, 2007; Vandermosten, Boets, Wouters, & Ghesquiere, 2012), however the direction of the correlation – positive versus negative – varies across studies and brain regions (Ben-Shachar et al., 2007; Deutsch et al., 2005; Klingberg et al., 2000; Lebel & Beaulieu, 2011; Niogi & McCandliss, 2006; Vandermosten et al., 2012; Yeatman et al., 2011), suggesting that the diffusion measurements in a given study may be influenced by different biological phenomena with distinct, and potentially opposing, relationships to reading (Yeatman, Dougherty, Ben-Shachar, & Wandell, 2012). Somewhat counter intuitively, a number of studies have reported *higher* FA in the commissural tracts of individuals with lower reading performance. This phenomenon has been attributed to a higher proportion of large diameter axons in this group(Dougherty et al., 2007), although it could also, theoretically, reflect reduced axonal dispersion (i.e., more highly skilled readers have more complex, less coherent, axonal architecture). Again, since DTI metrics lack biological specificity, there has been limited opportunity to test hypotheses about the underlying biological mechanisms that drive variation in reading skill.

The goals of the present work are threefold. First, we assess the scan-rescan reliability of biologically specific white matter indices derived from the WMTI and NODDI models in dMRI data collected in a group of children with varying ages (7-12 years) and varying reading levels (including children with dyslexia and typical readers). We show that the derived values in our sample are reliable and consistent with previously reported measurements. Second, we explore how decisions made in preprocessing affect model reliability. We find that image smoothing or de-noising is unnecessary when data are analyzed within individually defined white matter tracts using a robust measure, such as the median, to estimate values within a region of interest. Third, we examine individual differences in white matter maturation and reading skill, and demonstrate that these models can be used to test specific hypotheses about the correlation between reading skill and white matter diffusion properties. These results highlight the potential for biophysical modeling to enrich our understanding of the biological bases of cognitive development in health and disease.

## 2. Methods

### 2.1. Participants

Diffusion MRI and reading measures were collected for 55 children, ranging in age from 7 to 12 years. Each subject completed a series of reading tests, followed by an MRI scanning session. Subjects had a wide range of reading abilities, as assessed using the Woodcock-Johnson Basic Reading composite (untimed word and pseudo word reading accuracy): Age-normed scores ranged from 52 to 121, with a sample standard deviation of 14.084 (population mean=100, standard deviation=15). Of these subjects, 19 were used to assess scan re-scan reliability by collecting repeated dMRI measurements ranging from 2 to 8 weeks apart.

All participants were native English speakers with normal or corrected-to-normal vision and no history of neurological damage or psychiatric disorder. Subjects were screened using a mock scanner to assess comfort and ability to hold still during the MRI sessions. We obtained written consent from parents, and verbal assent from all child participants. All procedures, including recruitment, consent, and testing, followed the guidelines of the University of Washington Human Subjects Division and were reviewed and approved by the UW Institutional Review Board.

### 2.2. Diffusion MRI acquisition and pre-processing

All imaging data were acquired using a 3T Phillips Achieva scanner (Philips, Eindhoven, Netherlands) at the University of Washington Diagnostic Imaging Sciences Center (DISC) using a 32-channel head coil. An inflatable cap minimized head motion, and participants were continuously monitored through a closed circuit camera system.

Diffusion-weighted magnetic resonance imaging (dMRI) data were acquired at 2.0mm^3^ spatial resolution with full brain coverage. Each session consisted of two DWI scans, one with 32 non-collinear directions (b-value = 800 s/mm^2^), and a second with 64 non-collinear directions (b-value=2,000 s/mm^2^). Each of the DWI scans included 4 volumes without diffusion weighting (b-value=0), and the TE (85 ms) was held constant across scans. These acquisition values were chosen to optimally estimate both the NODDI and WMTI model parameters. In addition to these data, a scan with 6 non-diffusion-weighted volumes with a reversed phase encoding direction (posterior-anterior) was also collected to correct for EPI distortions due to inhomogeneities in the magnetic field using FSL’s *topup* tool(Andersson, Skare, & Ashburner, 2003). Additional pre-processing was carried out using tools in FSL for motion and eddy current correction(Andersson & Sotiropoulos, 2016). Data were manually checked for imaging artifacts and excessive dropped volumes. Given that subject motion can be especially problematic for the interpretation of group differences in dMRI data(Yendiki, Koldewyn, Kakunoori, Kanwisher, & Fischl, 2014), data sets with mean slice-by-slice displacement > 0.7mm were excluded from further analysis.

### 2.3 Biophysical model fitting and analysis

Diffusion metrics were estimated using the diffusion kurtosis model (Jensen, Helpern, Ramani, Lu, & Kaczynski, 2005), as implemented in DIPY(Garyfallidis et al., 2014). Axonal water fraction (AWF) and extra-axonal diffusivities were then estimated using the white matter tract integrity (WMTI) model (Fieremans et al., 2011; Fieremans et al., 2010), also implemented in DIPY. Following (Chung et al., 2018; Jensen et al., 2005), we restricted our analysis to voxels with high directional diffusion (fractional anisotropy > 0.3), to satisfy the modeling assumption of well-aligned fibers (Jensen et al., 2005). It is common practice to apply spatial smoothing to the DKI data prior to fitting the WMTI model, to reduce the influence of outliers. To minimize partial volume effects, we opted for block-wise non-local means de-noising (Coupe et al., 2008), implemented in DIPY. We also fit the model using raw DKI outputs, to evaluate the overall impact of a de-noising step in our analysis pipeline.

Intra-axonal volume fraction and orientation dispersion indices were estimated using the NODDI model (Zhang et al., 2012), implemented in Matlab. This provided maps of the intra-cellular volume fraction, orientation dispersion, and an isotropic (Viso) CSF fraction across the brain for each subject. The intra-cellular volume fraction (ICVF) is calculated by multiplying the intra-cellular volume fraction by 1 – Viso to obtain a quantity analogous to AWF provided by the WMTI model. The orientation dispersion index (ODI) is a mathematically independent metric that quantifies the coherence of fiber orientations, with a low value indicated aligned fibers.

Values from both modeling approaches were then mapped onto fiber tracts identified in each subject’s native space using the Automated Fiber Quantification (AFQ) software package(Yeatman, Dougherty, Myall, Wandell, & Feldman, 2012), after initial generation of a whole-brain connectome using probabilistic tractography (MRtrix 3.0)) (Tournier, Calamante, Gadian, & Connelly, 2004) (see https://github.com/yeatmanlab/afq/wiki for documentation). Fiber tracking was carried out on the aligned, distortion corrected, 64-direction (b-value = 2,000 s/mm^2^) datasets for each subject. After segmentation with AFQ, selected tracts were sampled into 100 evenly spaced nodes, spanning termination points at the gray-white matter boundary. Mean tracts values were estimated using the middle 60% of each tract, to minimize the influence of crossing fibers near cortical terminations, and to avoid potential partial volume effects at the white matter / gray matter border. AWF from the WMTI model and ICVF/ODI from the NODDI model were mapped onto each tract to create a ‘Tract Profile’. Here, we depart slightly from the methods described in (Yeatman, Dougherty, Myall, et al., 2012) and create these profiles by taking the median value at each node, rather than a distance weighted mean, to create a Tract Profile that is robust to outliers (afq.params.fiberWeighting = ‘median’). We compare results for median vs. weighted-mean (the default) Tract Profiles in **Figure 2**. Statistical analysis was carried out using software written in Matlab.

Data are available for download and visualization using AFQ-Browser (Yeatman, Richie-Halford, Smith, Keshavan, & Rokem, 2018) at: https://YeatmanLab.github.io/DevCogNeuro

## 3. Results

### 3.1 WMTI and NODDI parameters are reliable in children

We began by assessing scan-rescan reliability of the NODDI and WMTI parameters in the non-denoised data of a group of 19 children with repeated scanning sessions. As shown in **Figure 1**, all three of the relevant parameters were highly reliable: *r* = 0.79 median reliability for AWF (range = 0.52 to 0.93); *r* = 0.79 median reliability for ODI (range = 0.63 to 0.87); and *r* = 0.84 median reliability for ICVF (range = 0.53 to 0.92).

**Figure 1.**
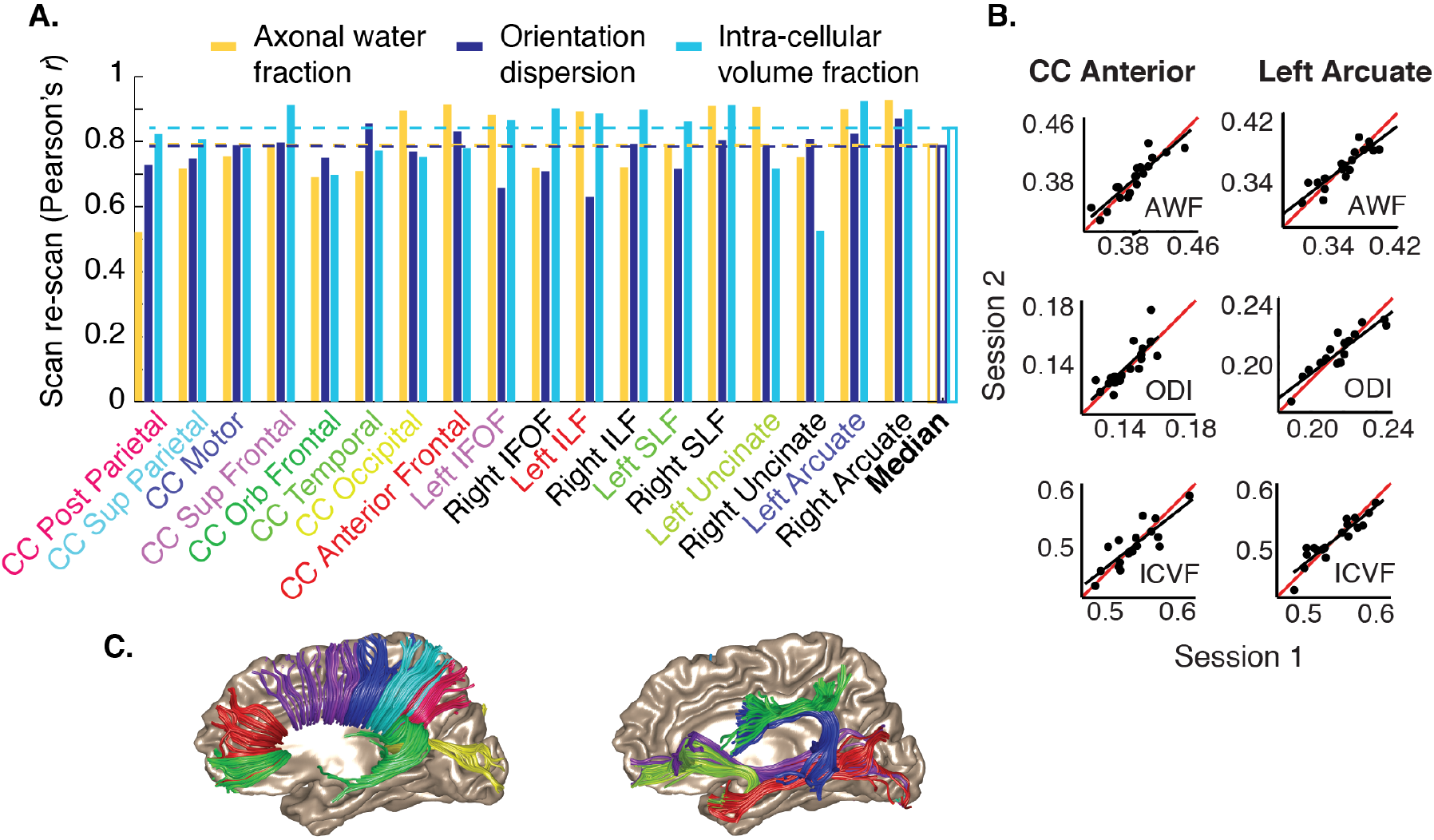
Parameters are reliable and within expected ranges. (A) Reliability (Pearson’s r) in a group of 19 subjects (ranging in age from 8-12 years) over repeated scanning sessions, separated by 2 to 8 weeks. Median reliability across all tracts (white bars, dashed lines) was greater than r = 0.75 for all three parameters. (B) Scatter plots show tract average axonal water fraction (AWF), intra-cellular volume fraction (ICVF), and orientation dispersion indices (ODI) for all subjects and two example tracts, estimated in Session 1 vs. Session 2. The identity line is shown in red, while the line of best fit is shown in black in each plot. (C) Example tract renderings. AWF, ICVF, and ODI values were mapped onto tracts crossing through 8 segments of the corpus callosum (left), and 10 additional intra-cortical association tracts (right). Color-coding of tract renderings corresponds to color-coding of tract names along the horizontal axis in (A).

The fitted values for AWF were consistent with previously reported estimates, ranging, for example, from 0.3 to 0.49 in the corpus callosum (Fieremans et al., 2011; Fieremans et al., 2010; Tang, Nyengaard, Pakkenberg, & Gundersen, 1997). Median AWF values were 0.31 for the posterior callosal tract (mean of 0.32, S.E.M across subjects of 0.0030). NODDI ICVF was consistently higher across tracts (as also noted previously, by (Jelescu et al., 2015)): Median ICVF was 0.54 in the posterior callosal tract (mean of 0.53, S.E.M across subjects of 0.0033). Median ODI was 0.15 for the callosal tract (mean 0.17, S.E.M. of 0.0043). This latter value is consistent with previous estimates based on dMRI and histology (Mollink et al., 2017). As expected, ODI values were slightly higher (0.2-0.3) for the association tracts.

### 3.2 Anatomically informed robust averaging substitutes for image de-noising

A low signal-to-noise ratio and presence outliers are of particular concern in developmental datasets since the time constraints associated with scanning children often precludes repeated scans within a session, and since these data sets are vulnerable to artifacts from motion and other factors. De-noising is a common additional pre-processing step prior to WMTI model fitting and it is thought to be important for improving the SNR of the data and reducing the influence of outliers in the DKI fits (Jelescu et al.; Veraart et al.). However, researchers are often faced with a challenge when trying to decide on the necessary and optimal de-noising steps for their data. To analyze the effects of de-noising on scan-rescan reliability, we preprocessed our data using a non-local means filtering approach with commonly used parameter settings (as described in **Section 2.3**), and without de-noising. As shown in **Figure 2**, use of de-noising filter size had a minimal influence on the results. De-noising produced highly reliable estimates, but the reliability was not higher than the non-denoised data. Given the trend for the median reliability to be slightly higher without de-noising, we opted to carry out the rest of our analyses using the non-denoised data set.

While this result might seem counter-intuitive – de-noising does not improve the scan-rescan reliability of parameter estimates – it makes sense in the context of a tract-based analysis (i.e., tractometry) where values are averaged over anatomically defined regions of interest. By using AFQ to extract median values from each white matter tract, we were able to alleviate the need for a de-noising step, since median values are inherently robust to outliers. For comparison, reliabilities are plotted for the non-denoised data sampling the core of the white matter using a using a distance weighted means approach (Yeatman, Dougherty, Myall, et al., 2012). Reliability is substantially better when values are calculated by taking the median, rather than the weighted-mean, of voxels in a tract, because outlier voxels with extreme values do not bias the median.

**Figure 2.**
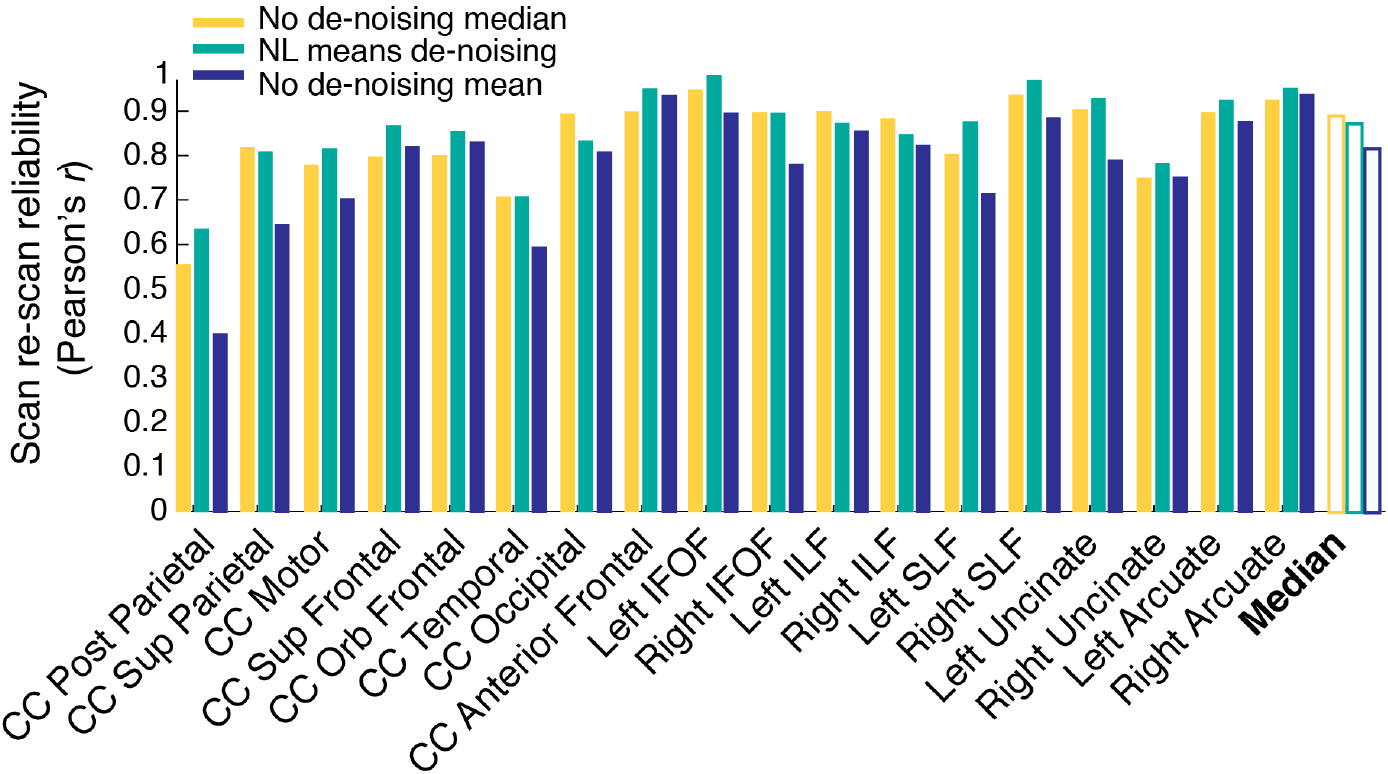
De-noising has a small effect on parameters derived from the WMTI model. Reliability (Pearson’s r) for each tract, using non-local means filtering for image de-noising (see **section 2.3** for details).

### 3.3 Developmental changes in white matter biology

Developmental changes in mean diffusivity and fractional anisotropy (FA) have been described throughout the white matter (Lebel & Beaulieu, 2011), and there has been recent interest in elaborating these results using biophysical modeling (Chang et al., 2015; Genc et al., 2017; Mah et al., 2017). Here, we began by examining developmental effects in the posterior callosal connections. We chose this as our target for three reasons. First, the high coherence of axons within the posterior callosal connections should allow for the most accurate estimates of axon properties based on the WMTI model, since the model is less interpretable in regions with complex fiber geometry. Second, histological studies in non-human primates (Hopkins & Phillips, 2010) and structural MRI studies in humans (Giedd et al., 1999; Giedd et al., 1996; Kim et al., 2007) indicate that the corpus callosum continues to develop and myelinate throughout childhood and into adolescence, and diffusion properties show large maturational effects within the age range of our sample (McLaughlin et al., 2007; Snook, Paulson, Roy, Phillips, & Beaulieu, 2005). Finally, the posterior callosum plays a particularly important role in the literature relating DTI measures to reading skills (Dougherty et al., 2007).

As shown in **Figure 3a**, ICVF and AWF both increase as a function of age in this tract, while ODI declines. Thus, previously reported variation in FA likely reflects at least two distinct phenomena: Both the increase in AWF/ICVF and the decrease in ODI would contribute to an increase in FA during childhood.

We then carried out an exploratory analysis including all of the commissural and association tracts. We summarize the results of that analysis in **Figure 3b**. In general, ODI decreased as a function of age, while ICVF and AWF increased, and effect sizes varied across tracts.

**Figure 3.**
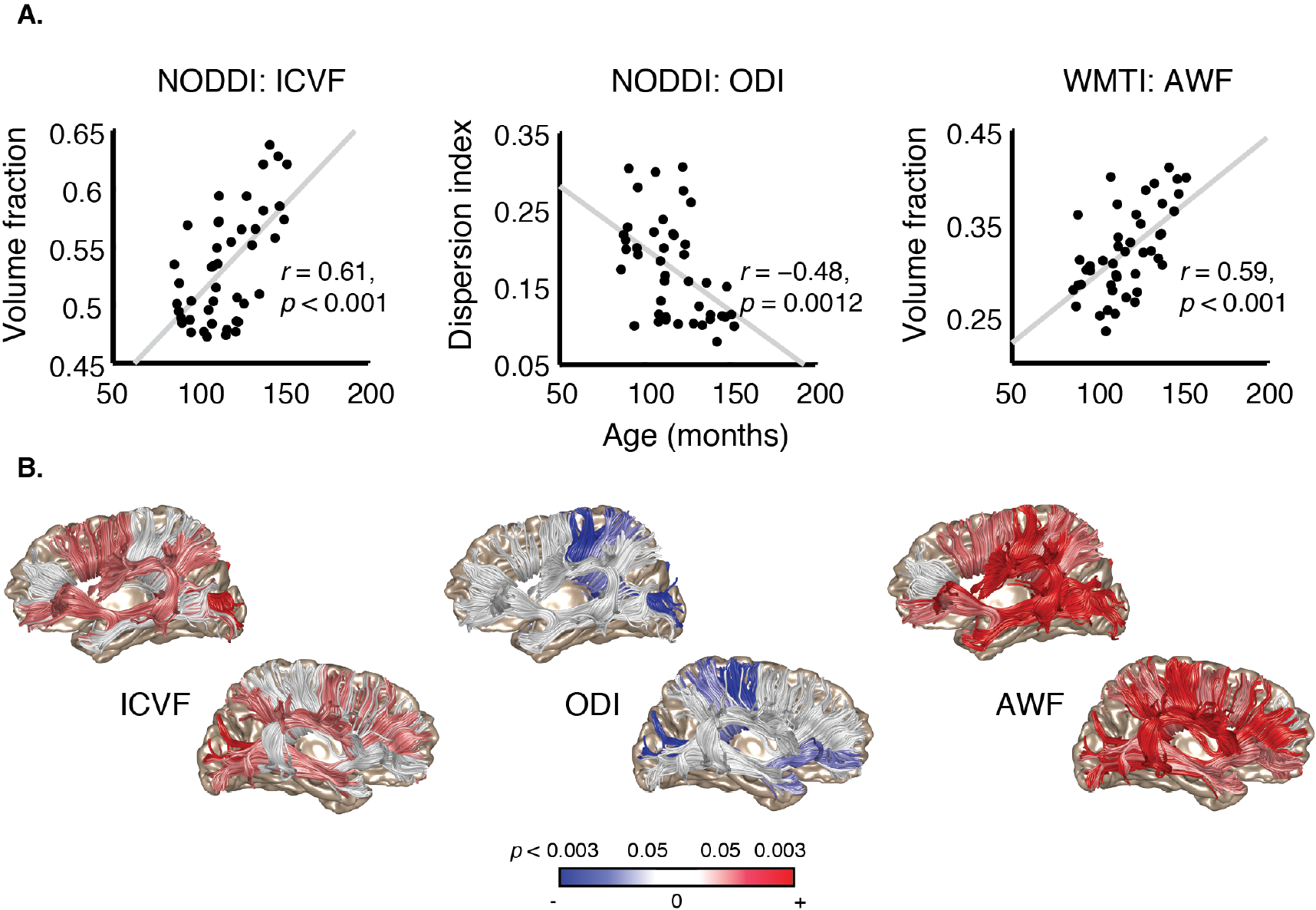
Developmental changes in the callosal connections and association tracts. (A) Tract average AWF, ODI, and ICVF values plotted as a function of age (in months) for the posterior callosal connections. (B) Anatomical rendering, with tracts showing significant (p < 0.05, Bonferroni corrected) age related variation in ICVF, ODI, or AWF highlighted in red (positive correlation) or blue (negative correlation). Tracts rendered in less saturated red and blue show moderate age-related variation (p < 0.05, uncorrected).

### 3.4 Biological underpinnings of reading-diffusion correlations

The structure of the posterior corpus callosum has previously been shown to differ in both children and adults with dyslexia (Duara et al., 1991; Rumsey et al., 1996; von Plessen et al., 2002), and several studies have reported correlations between reading-related skills and diffusion properties within posterior callosal regions (Dougherty et al., 2007; Frye et al., 2008; Hasan et al., 2012; Odegard, Farris, Ring, McColl, & Black, 2009). In the diffusion literature, higher radial diffusivity and lower FA for this tract have been associated with higher reading proficiency. These effects are somewhat counter intuitive, given that they would suggest reduced density of inter-hemispheric connections in strong readers, but they have been linked to the hypothesis that reading-related functions are not as strongly left-lateralized in struggling readers (Finn et al.; Galaburda, Aboitiz, Rosen, & Sherman; Galaburda, Sherman, Rosen, Aboitiz, & Geschwind, 1985; Rumsey et al.). A higher proportion of large diameter axons in this region in struggling readers could account for this effect, as hypothesized by (Ben-Shachar et al., 2007), although reduced axonal dispersion could also account for the higher FA.

**Figure 4.**
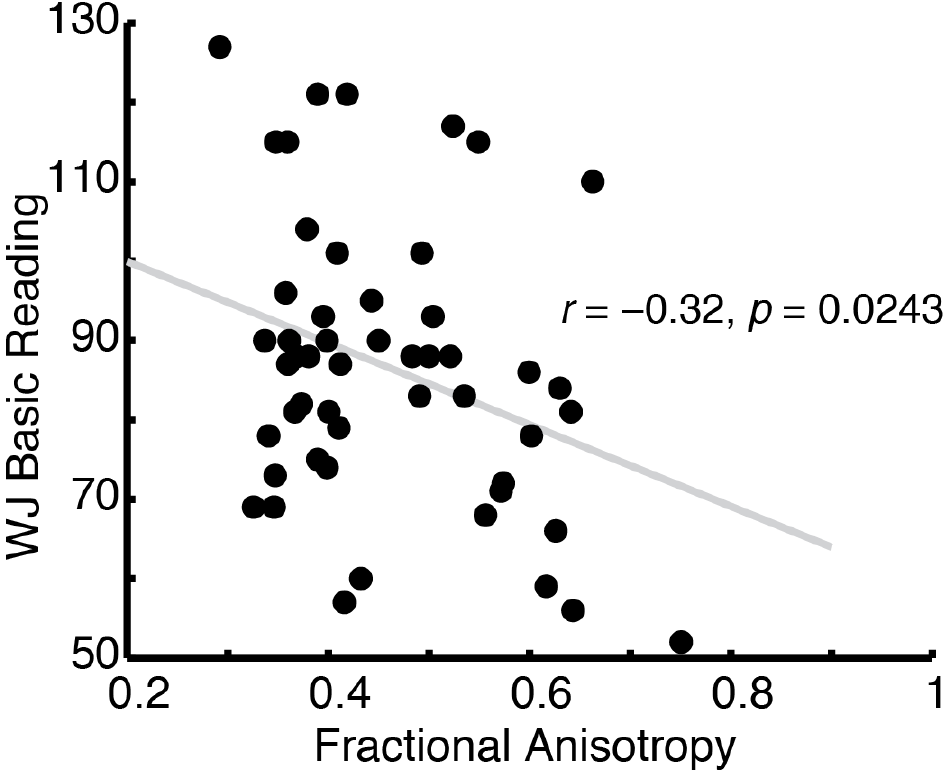
Fractional anisotropy (FA) in the posterior callosum is negatively correlated with reading skill. Woodcock-Johnson Basic Reading scores are plotted as a function of tract average FA values from the posterior callosal connections. Replicating previous studies, we find a negative relationship between FA and reading skill within this region: Individuals with higher FA are generally worse readers.

Here, we replicated the previously reported negative correlation between reading skills and FA in the posterior callosal tract (**Figure 4**). We then examined the relationship between NODDI ICVF, NODDI ODI, and WMTI AWF values and reading skills in the same region. As shown in **Figure 5**, ICVF and AWF both significantly correlate with reading skill (Pearson’s *r*), while ODI does not. Although we report standardized reading scores, it is possible that confounding effects of age might exist since, for example, older struggling readers often have lower standardized reading scores (i.e., lower scores relative to their peers). To control for a possible confounding effects of age, given the age dependence of the NODDI and WMTI parameters, we fit a linear model predicting reading scores from each parameter with age included as a covariate. In this analysis, NODDI ICVF accounted for a significant proportion of the variance in reading skill (*F*(1,41) = 5.33, *p* = 0.026). Meanwhile, NODDI ODI was not a significant predictor of reading skill, over and above age: *F*(1,41) = 1.33, *p* = 0.25. In line with this result, AWF also predicted reading skill, over and above age: *F*(1,43) = 6.14, *p* = 0.017.

Finally, we carried out an exploratory analysis of each additional tract in the data set. As shown in **Figure 5**, only ICVF in the anterior callosal tract and ODI in the right arcuate significantly predicted reading skill, after controlling for age. Thus, the correlation between axon properties and reading skill was relatively specific to the posterior callosum.

**Figure 5.**
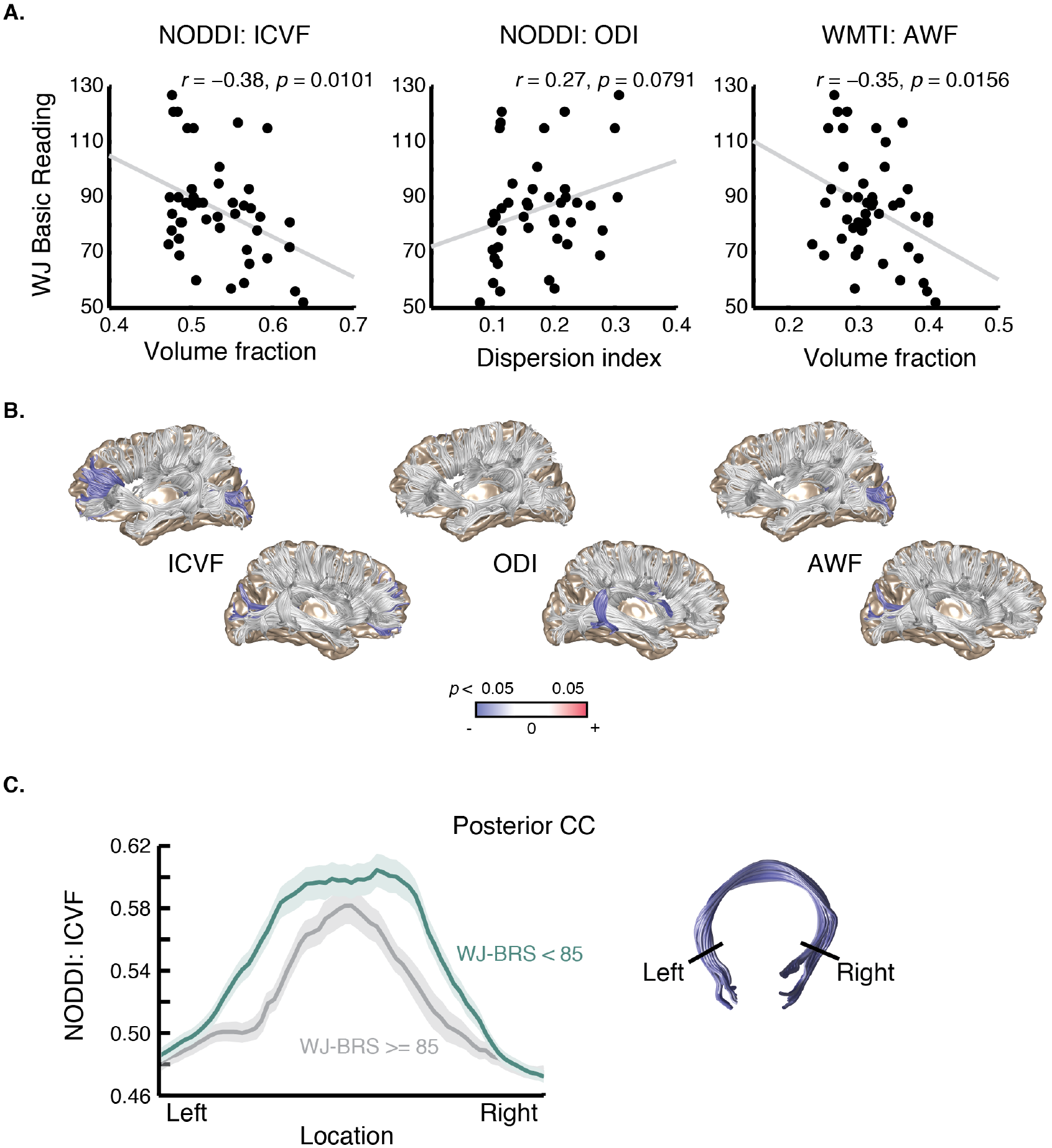
Indices of tissue density in the posterior callosal connections correlate with reading skill. (A) Scatter plot showing Basic Reading as a function of NODDI ICVF, ODI, and WMTI AWF values. Insets show tract profiles for skilled vs. struggling readers for each parameter (mean +/− 1 standard error). (B) Anatomical rendering, with tracts showing significant (p < 0.05) reading related variation, after controlling for age, ICVF, ODI, or AWF highlighted in red. (C) Tract profile along the posterior callosal connections showing mean AVF for skilled readers (WJ Basic Reading score at or above 85) and struggling readers (WJ Basic Reading score below 85). The x-axis spans the middle 60% of each tract the where it was clipped prior to analysis (corresponding to black boundary lines in the example anatomical renderings, at right). Shading represents one standard error of the mean.

## 4. Discussion

Here we use two popular biophysical modeling techniques, “White Matter Tract Integrity” or WMTI (Fieremans et al., 2011; Fieremans et al., 2010) and “Neurite Orientation Dispersion and Density Imaging or NODDI, to examine white matter properties related to development and individual differences in reading skill in a group of grade-school aged children. Both models provided metrics that were reliable and within the expected range based on previous estimates from histology and *in vivo* microscopy (reviewed in (Jelescu & Budde, 2017)). Additionally, we were able to fit the WMTI model with minimal image pre-processing by using the AFQ tractometry pipeline to generate anatomically informed, robust estimates from the data. We then used the estimated parameters from each model to examine age- and reading skill-related variation in white matter properties, highlighting the utility of these models for testing specific hypotheses about the biology of the white matter in relation to cognitive development.

Model-estimated axonal water fraction (AWF) and intra-cellular volume fraction (ICVF) increased linearly with age in a large collection of anatomical tracts. ICVF increased with age bilaterally in the inferior longitudinal fasciculus and the posterior and superior frontal callosal tracts. ICVF also increased in the left arcuate fasciculus and inferior frontal fasciculus, and in the right superior longitudinal fasciculus. AWF increased in nearly every tract measured. Meanwhile, ODI decreased in a few callosal tracts – the posterior, superior parietal and motor tracts – and in the right uncinate, but was otherwise stable. Together, these factors should contribute to a widespread developmental increase in FA, as reported elsewhere (Lebel & Beaulieu, 2011). A recent study examining FA and NODDI metrics within the same age group (7-12) showed a similar pattern: On average, cortical tracts show relatively stable ODI during the first decade of life, with a gradual increase that accelerates over the lifespan, while neurite density increases sharply (Chang et al., 2015), suggesting that developmental changes in white matter diffusion primarily reflect increases in axonal diameter and myelination.

The successful application of these models in the context of development opens the possibility that they can also be applied to examine functionally relevant features of the white matter. For instance, axon caliber is intrinsically linked to signal conduction velocity and firing rate, and hence theoretically linked to the maximal rate of information encoded by a given neuron (Perge, Niven, Mugnaini, Balasubramanian, & Sterling, 2012). Differences in white matter diffusion that correlate with cognitive skills are often interpreted as reflecting enhanced signaling efficiency, although metrics like FA and MD are too far divorced from axonal properties to allow a direct test of these ideas. Here, we take an example from the reading literature and test whether biologically based metrics can be applied to examine individual differences in single-word reading accuracy. Diffusion anisotropy of the posterior callosal tract has previously been linked to reading skill (Alhamud et al., 2012; Dougherty et al., 2007; Frye et al., 2008; Odegard et al., 2009), and it has been hypothesized that this reflects increased inter-hemispheric connectivity, mediated by denser axon packing, and/or larger caliber axons, in struggling readers (Ben-Shachar et al., 2007). Using biologically based modeling, we were able to test this hypothesis: We found that axonal water fraction and intra-cellular volume fraction of the posterior callosal tract correlate with reading skill, while orientation dispersion does not. This suggests the elevated FA in struggling readers indeed reflects larger caliber axons and/or denser axonal packing that could support a greater capacity for inter-hemispheric signaling. This anatomical observation is consistent with functional MRI studies showing that activation patterns in a variety of language and reading tasks are less left lateralized in the dyslexic brain.

Although AWF and ICVF both increased over development in the posterior callosal connections, these properties also differed as a function of reading skill, independent of age. Differences in axonal properties within this region may therefore emerge early in life and influence subsequent development throughout the reading circuitry, even as the callosal connections themselves continue to mature. In line with this idea, the posterior callosal tract was remarkably stable within subjects during an 8-week, intensive reading intervention that prompted large changes in diffusion properties throughout a collection of cortical association and projection tracts(Huber, 2018). Indeed the callosum was one of just a few tracts that did not change during the intensive reading skills training program. Thus, anatomical differences that are stabilized prior to age 7 (the youngest individuals included in our sample) in the posterior callosal tract may ultimately shape reading development, while other portions of the reading network remain amenable to change, perhaps reflecting compensatory mechanisms that can emerge with educational intervention (Barquero, Davis, & Cutting, 2014; Eden et al., 2004; Hoeft et al., 2011; Shaywitz et al., 2004). Future work linking white matter biology to functional responses within the reading circuitry should help to build a more nuanced view of the computations involved, and the ways in which these circuits develop and adapt to experience.

## 5. Conclusions

Biophysical modeling offers a bridge between diffusion MRI and histology, allowing us to test specific, biologically based hypotheses about cognitive development. If data processing steps are taken to ensure reliable parameter estimates, this approach holds promise for adding nuance to our understanding of the biological changes that occur in human white matter over development and for revealing the computational mechanisms associated with individual differences in cognition.

## Acknowledgements

This work was funded by NSF/BSF BCS #1551330 to JDY. The International Neuroinformatics Coordinating Facility (INCF) provided support through a project and travel grant to RNH. AR was funded through a grant from the Gordon & Betty Moore Foundation and the Alfred P. Sloan Foundation to the University of Washington eScience Institute.

